# AdhE spirosome length in enterohaemorrhagic *Escherichia coli* is correlated with enzymatic directionality and is perturbed by salicylidene acylhydrazide binding

**DOI:** 10.1101/2024.01.25.577206

**Authors:** Ester Serrano, Arwen I. I. Tyler, Mostafa Soroor, Iris Floria, Nikil Kapur, Andrew J. Roe, Olwyn Byron

## Abstract

Antibiotics are contraindicated for the treatment of infection enterohemorrhagic *Escherichia coli* (EHEC), a human pathogen that causes diarrhea or hemorrhagic colitis in humans which can progress to hemolytic uremic syndrome (HUS). As an alternative to the use of antibiotics, previous studies developed the salicylidene acylhydrazides (SA), a family of anti-virulence compounds capable of blocking expression of the type three secretion system (T3SS), thereby reducing bacterial infections. Here we validate AdhE as the anti-virulence protein target of the SA compound ME0054. AdhE is a bidirectional enzyme able to catalyse the conversion of acetyl-CoA to ethanol and *vice versa*. AdhE oligomerises forming helicoidal filaments heterogeneous in length called spirosomes. In this work, we show that it is possible to partially fractionate AdhE spirosomes using size exclusion chromatography (SEC) and to characterise the spirosome oligomers present in each fraction with biophysical techniques such as small angle X-ray scattering (SAXS) and sedimentation velocity analytical ultracentrifugation (SV-AUC). Also, we observe that short spirosomes are more efficient in the reverse reaction whereas the spirosome length has no impact on the forward reaction. Therefore, for first time, we reveal that AdhE spirosome formation is necessary to regulate the direction of its enzymatic reactions. In addition, we show that ME0054 disrupts AdhE spirosomes, thereby enhancing the conversion of ethanol to acetyl-CoA. Importantly, SV-AUC data show that ME0054 binds to the AdhE filaments. Finally, time-resolved (TR) SAXS allowed us to follow the kinetics of spirosome disruption produced by ME0054, confirming its effectiveness at biologically relevant temperatures and timescales.

**SIGNIFICANCE STATEMENT:** There is an urgent need to develop alternative strategies to combat bacterial infections. Salicylidene acylhydrazides (SA) are able to reduce expression of the bacterial type three secretion system (T3SS), used by many pathogens to manipulate host eukaryotic cells, including our pathogen of interest: enterohaemorrhagic *E. coli* (EHEC). The mechanism underpinning these compounds is a mystery. Here we show how the SA compound ME0054, by disrupting AdhE spirosomes, enhances metabolic conversion of ethanol to acetyl-CoA. This finding is consistent with the phenotype observed in an EHEC AdhE mutant: alterations in acetate levels and changes in T3SS expression. Our work establishes a crucial mechanistic link between the binding of the SA compound to a key target protein and changes in bacterial metabolism.

## INTRODUCTION

Addressing bacterial infections has grown progressively challenging in light of the rising prevalence of antibiotic-resistant strains and a decline in the creation of innovative antibacterial agents (1). Presently, the majority of antibiotics operate by inhibiting enzymes crucial for pathogen survival. For instance, β-lactams and aminoglycosides target bacterial cell wall biosynthesis and translation, respectively. As these processes are essential for growth, the selective pressure imposed by antibiotics is strong, thereby increasing the forces driving antibiotic resistance. The identification of novel targets that are not essential for survival is therefore an active area of research.

The so called “anti-virulence” (AV) approach is one that specifically targets virulence factors used by bacterial pathogens to facilitate the infection process (2). AV compounds designed or screened to target important virulence traits such as quorum sensing systems, adhesins, and secretion systems have been tested, however the development of these compounds is still in the early stages. AV approaches also have advantages when the use of traditional antibiotics is not appropriate. For example, the clinical symptoms associated with enterohaemorrhagic *Escherichia coli* (EHEC) infections have been shown to increase in severity following administration of certain antibiotics (3). This is a result of the release of Shiga toxin following bacterial lysis (4). Therefore, by targeting virulence, these negative side effects may be mitigated.

One of the most extensively studied group of AV compounds is the salicylidene acylhydrazides (SA), a class of inhibitors that were identified from a chemical screen of 9400 compounds carried out by Kauppi *et al*. at the University of Umeå (5). The screen was performed on *Yersinia pseudotuberculosis* expressing a *yopE*-luciferase transcriptional fusion. YopE is a secreted effector protein; therefore, a decrease in luciferase activity from the promoter was correlated with reduced expression of the type three secretion system (T3SS). Compounds of this family have subsequently been shown to be capable of inhibiting the expression of the T3SS of numerous additional pathogens including *Chlamydia trachomatis, Salmonella* Typhimurium, *Shigella flexneri* and EHEC (6).

The precise mechanism by which the SA compounds block expression of the T3SS is not established. Surprisingly, none of the structural proteins involved in the assembly of the T3SS was found to be a target of the SA compounds. Rather, it was demonstrated that multiple, conserved, bacterial proteins were bound by the SA compounds (7). Of these, the most interesting was AdhE, a bifunctional aldehyde-alcohol dehydrogenase able to catalyse the bidirectional conversion of acetyl-CoA to ethanol (8-11). Deletion of the gene (*adhE*) encoding AdhE resulted in a strong suppression of the T3SS and elevated levels of extracellular acetate. This phenotype is similar to that observed when this strain is exposed to SA compounds (8). However, how the SA compounds cause this phenotype was unresolved and has spurred us to better understand the structure and function of AdhE.

AdhE oligomerises both *in vivo* and *in vitro*, forming long filaments, heterogeneous in length, called spirosomes (12). The cryo-electron microscopy (cryo-EM) structure of the spirosome from *E. coli* has been reported and mutations disrupting the helical structure revealed that spirosome formation is critical for AdhE activity (9). However, why AdhE forms spirosomes heterogeneous in length is still a mystery. We envisioned at least two hypotheses to explain why AdhE forms spirosomes with different lengths: (a) longer spirosomes are enzymatically more efficient or (b) the spirosome length drives the direction of the enzymatic reaction. To test these hypotheses, in this work: (i) AdhE spirosomes were fractionated by length, (ii) the oligomeric species present in each fraction were analysed by small angle X-ray scattering (SAXS) and analytical ultracentrifugation (AUC), (iii) fractions containing a main population of long or short spirosomes were selected and (iv) their efficiencies were compared both in the forward and the reverse direction of the enzymatic reaction. The effect of the SA compound ME0054 (7, 13) was then tested for effects on spirosome structure and enzymatic efficiency to provide new insights into how the compounds perturb AdhE structure and function.

## RESULTS

### AdhE spirosomes are partially fractionated by size exclusion chromatography (SEC)

AdhE with a 6-His tag fusion at the N-terminal was purified using a Ni-NTA column. As previously described (9, 11), we observed by negative stain electron microscopy (EM; Fig. 1*A*) how purified AdhE formed spirosomes heterogeneous in length. Size exclusion chromatography (SEC) was used to fractionate the AdhE spirosomes by hydrodynamic volume. AdhE eluted as a single peak, so to try to obtain samples homogeneous in spirosomes of different lengths, six 2 ml fractions (termed F12, F14, F16, F18, F21 and F24) were collected across the elution peak (Fig. 1*B*). These fractions were analysed by sedimentation velocity (SV) AUC which revealed that none of the fractions was homogeneous, all c(s) profiles comprising multiple peaks (Fig. 1*C*). This indicates that the AdhE spirosomes were only partially fractionated by SEC. The c(s) profiles for later eluting fractions are increasingly dominated by lower sedimentation coefficient species, consistent with a reduction in spirosome oligomeric state.

**Figure 1.**
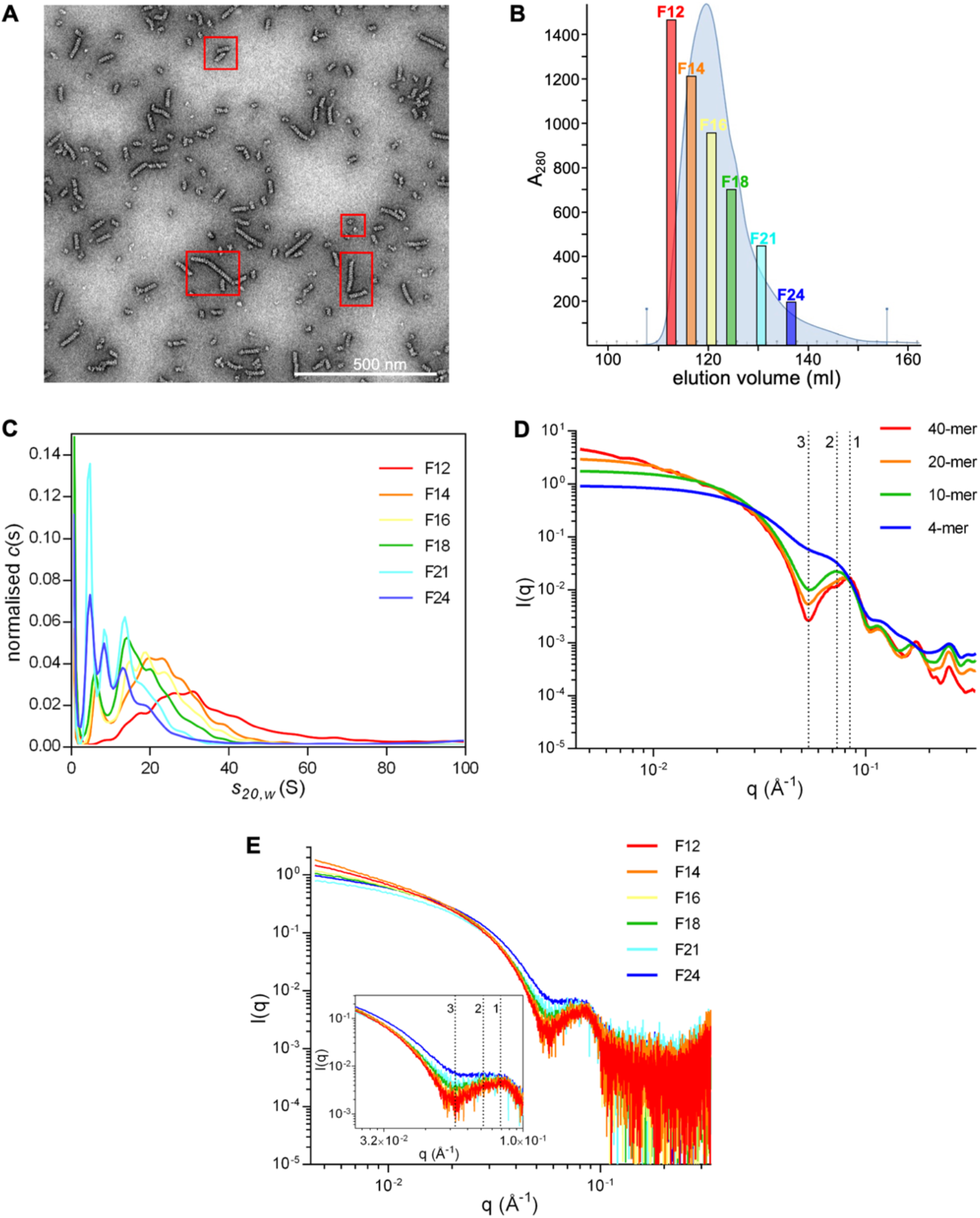
Characterisation of AdhE spirosome fractionation. **(A)** Unfractionated AdhE visualised by negative stain electron microscopy (EM). Different length spirosomes are highlighted by red squares. **(B)** Purified AdhE was fractionated by size exclusion chromatography (SEC) and six representative fractions were selected from the elution peak. **(C)** Sedimentation velocity analytical ultracentrifugation (SV-AUC) analysis of the six representative AdhE fractions. Small angle X-ray scattering (SAXS) profiles **(D)** computed with FoXS server [11,12] for four spirosome models of different lengths and **(E)** obtained experimentally in batch mode from the six representative AdhE fractions. The positions of three conserved features are indicated at q = 0.086 Å^−1^ or q = 0.083 Å^−1^ (feature 1, in computed and experimental profiles, respectively), q = 0.072 Å^−1^ (feature 2) and q = 0.052 Å^−1^ (feature 3).

Previous work (9) correlated a conserved feature in AdhE spirosome small angle X-ray scattering (SAXS) profiles at q = 0.086 Å^−1^ with the helical pitch of the spirosome cryo-EM structure. We hypothesised here that the loss of helical repeats when the spirosome length decreases will be reflected in a change in this conserved feature. To test this, SAXS profiles were computed with the FoXS server (14, 15) for linear spirosome models of different lengths (4-, 10-, 20- and 40-mer; Fig. 1*D*), constructed from the atomic resolution structure of AdhE (PDB ID 6AHC (9)). As expected, a dramatic shift in the reciprocal space position (q) of the conserved feature was observed when the spirosome model length decreased from 40-mer (q = 0.083 Å^−1^) to 4-mer (q = 0.072 Å^−1^) (Fig. 1*D*), supporting our hypothesis. The marginal difference in q for this feature in the SAXS profiles computed for AdhE spirosome models (q = 0.086 Å^−1^) compared with that observed experimentally (q = 0.083 Å^−1^) most probably arises from sample heterogeneity, the presence of small spirosomes in fractions otherwise dominated by larger filaments shifting the position of this feature slightly. The loss of helical repeats was also reflected in a gradual reduction in the slope at low angles (q < 0.025 Å^−1^), consistent with a lower radius of gyration corresponding to shorter spirosomes. Interestingly, a very pronounced trough at position q = 0.052 Å^−1^ diminished progressively with the reduction in length, becoming a new indicator within the SAXS profile for the analysis of spirosome length (Fig. 1*D*).

SAXS data for the six representative AdhE fractions were obtained in batch mode. Comparison of the experimental data (Fig. 1*E*) with the profiles computed for the four spirosome models (Fig. 1*D*) confirms that the earliest eluting faction (F12) is dominated by large oligomers whilst fractions collected thereafter comprise increasingly shorter spirosomes. To estimate the average spirosome length in each fraction, the experimental data were fitted with profiles computed by the FoXS server (14, 15) for a wider series of oligomeric models. The oligomeric states that provided the best fits for each fraction are reported in Table S1, reflecting a decrease in dominant spirosome length from F12 to F24 of 40-mer to 14-mer. In heterogeneous samples, SAXS profiles are dominated by the scattering from bigger species. Thus, although later eluting fractions, such as F24, are shown by SV-AUC to be dominated by smaller species (Fig. 1*C*), the presence of larger oligomers will result in elevated values of χ^2^ for the fit to the experimental data by the models, as observed in Fig. S2, Table S1.

### Spirosome length drives the direction of the AdhE enzymatic reaction

AdhE is a bidirectional enzyme (8-11) able to both convert acetyl-CoA to ethanol (forward reaction) and ethanol to acetyl-CoA (reverse reaction) (Fig. 2*A*). To explore whether spirosome length could be related to the efficiency of the enzymatic reaction, the enzymatic activity of F12 and F24 (dominated by long or short spirosomes, respectively) was tested in the forward and reverse reactions. To measure the forward reaction, F12 or F24 were incubated with acetyl-CoA in the presence of NADH and the activity measured by monitoring the consumption of NADH at 340 nm. No difference in activity was observed between F12 and F24 suggesting that the spirosome length has no impact on AdhE activity in the forward reaction (Fig. 2*B*). To measure the reverse reaction, F12 or F24 were incubated with ethanol, NAD^+^ and CoASH, and the activity measured by monitoring the production of NADH at 340 nm. Surprisingly, in the reverse reaction, the fraction dominated by short spirosomes (F24) was much more efficient than that dominated by long spirosomes (F12) (Fig. 2*C*). Taken together these results suggest that AdhE needs to oligomerise to form spirosomes in order to balance the direction of the enzymatic reaction.

**Figure 2.**
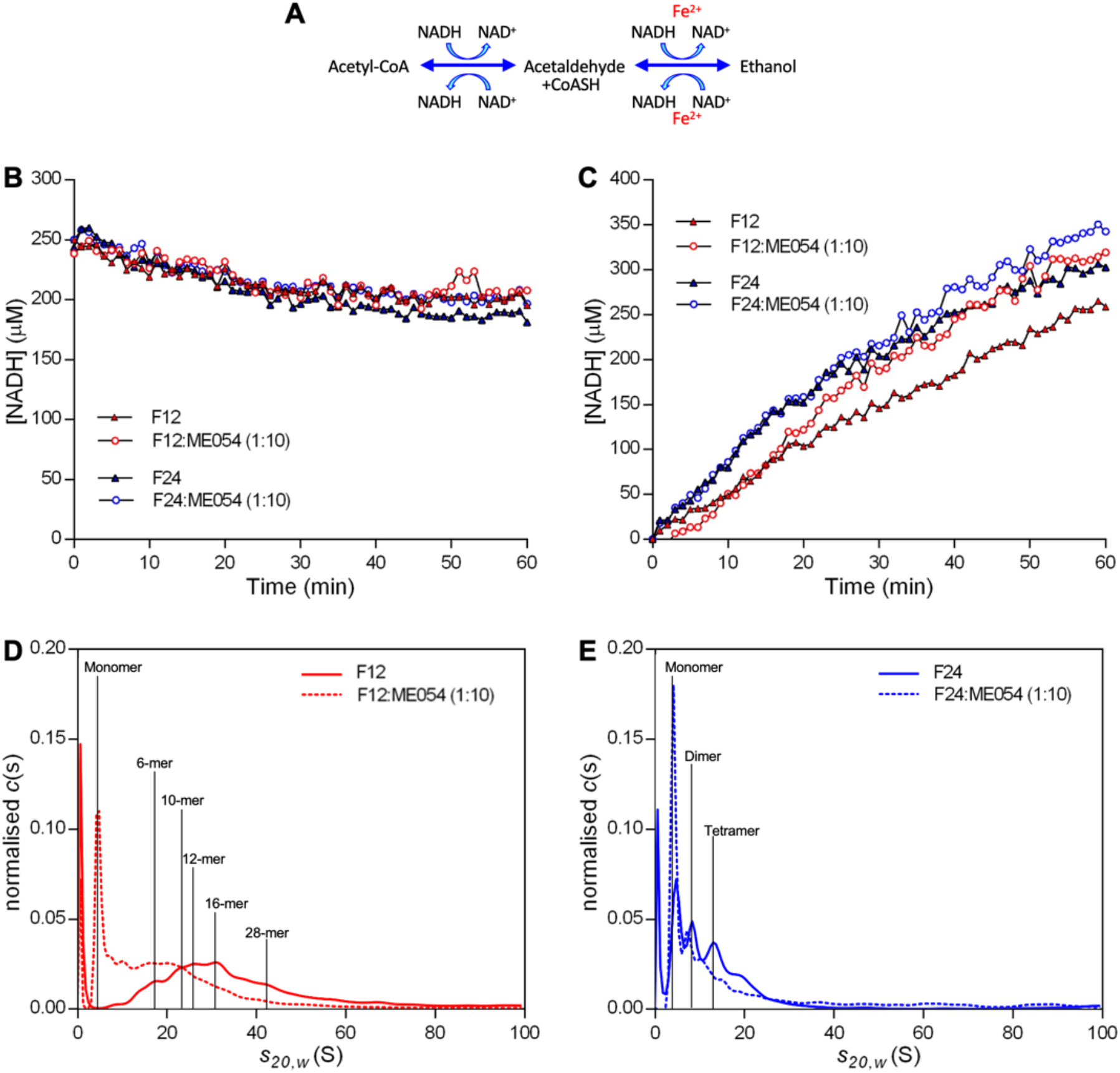
AdhE spirosome length drives the direction of the enzymatic reaction. **(A)** Schematic representation of the enzymatic reactions catalysed by AdhE. Enzymatic assays of AdhE fractions containing a main population of long (F12, red symbols) or short (F24, blue symbols) spirosomes in the presence (open circles) or absence (filled triangles) of ME0054 (1:10 molar ratio) during the forward **(B)** or reverse **(C)** reactions. ME0054 disrupted AdhE spirosomes enhancing the reverse reaction where short spirosomes are more efficient. c(s) analysis of SV-AUC data for **(D)** F12 and **(E)** F24 +/- 1 h incubation with ME0054 at 37°C.

### ME0054 binds to AdhE spirosomes and disrupts them in a concentration dependent manner

Deletion of the *adhE* gene in EHEC results in a marked reduction in T3SS activity and an increase in flagella production (8), both of which are phenotypes observed when this strain is exposed to the SA compounds (16). Previous studies proposed the AdhE protein as a target of the SA compounds (8) in EHEC. To examine whether the SA compounds affect the enzymatic activity of AdhE, F12 and F24 were incubated with the SA compound ME0054 at a molar ratio of 1:10 (AdhE:ME0054) during the forward and reverse reactions (Fig. 2*B, C*). No impact on AdhE activity was observed in the presence of ME0054 during the forward reaction (Fig. 2*B*). However, in the reverse reaction, the efficiency of long spirosomes (F12) dramatically increased after 17 min in the presence of ME0054, reaching a level comparable to that attained by short spirosomes (F24; Fig. 2*C*). In contrast, the activity of F24 (dominated by shorter spirosomes) only slightly increased in the presence of ME0054 (Fig. 2*C*). These data suggest that ME0054 disrupts AdhE spirosomes, transforming the oligomeric profile of F12 to something resembling F24.

To explore this idea, F12 and F24 were incubated with ME0054 for 1 h at 37°C at a molar ratio of 1:10 (AdhE:ME0054) and analysed by SV-AUC (Fig. 2*D, E*). The size distribution for F12 clearly shifted towards lower sedimentation coefficients in the presence of ME0054, with the dramatic appearance of a peak at s_20,w_ = 5.2 S, confirming that the compound disrupts long (F12) spirosomes (Fig. 2*D*). When short spirosomes (F24) were incubated with ME0054 only a slight shift in the size distribution (towards lower sedimentation coefficients) was observed (Fig. 2*E*), reflecting the disruption in the few larger spirosomes present in this fraction. To further this analysis, we wanted to determine the spirosome oligomeric state in these fractions before and after the addition of ME0054. Linear spirosome models between monomer and 54-mer were generated based on the atomic structure of AdhE (PDB ID 6AHC (9)). AtoB (17) was used to compute grid-based bead models for which hydrodynamic calculations were undertaken using SOMO (18, 19) and for each oligomer the corresponding sedimentation coefficient was plotted (Fig. S1). Based on this, it was possible to assign the probable oligomeric states across the SV-AUC size distribution analysis (Fig. 2*D, E*). AdhE F12 contained a main population of 12-16-mer spirosomes which were disrupted by ME0054 to 4-10-mers, with the appearance of a significant population of monomers (Fig. 2*D*). When F24 was incubated with ME0054, the number of tetramers was reduced whilst the population of monomers increased (Fig. 2*E*).

To further investigate the disruption of AdhE spirosomes by ME0054, F12 was incubated with ME0054 at different molar ratios for 1 h at 37°C for visualisation of the effect by negative stain EM (Fig. 3*A*). The lengths of at least 100 molecules per condition were quantified (Fig. 3*B*), concluding that the effect of ME0054 on AdhE spirosomes is concentration dependent. To confirm that small molecules were not missed during the quantification due to the limits of the microscope magnification, the same conditions were tested by SV-AUC at wavelengths of 280 nm (Fig. 3*C*) and 330 nm (Fig. 3*D*) to observe AdhE and ME0054, respectively. The SV-AUC data recorded at 280 nm support the conclusion that ME0054 disrupts AdhE spirosomes in a concentration dependent manner, since the position of the main peak in each distribution shifts towards lower sedimentation coefficients as the compound concentration increases. The SV-AUC data recorded at 330 nm show that the compound binds to AdhE spirosomes, because at all molar ratios ME0054 co-sedimented with both long and short spirosomes (Fig. 3*D*).

**Figure 3.**
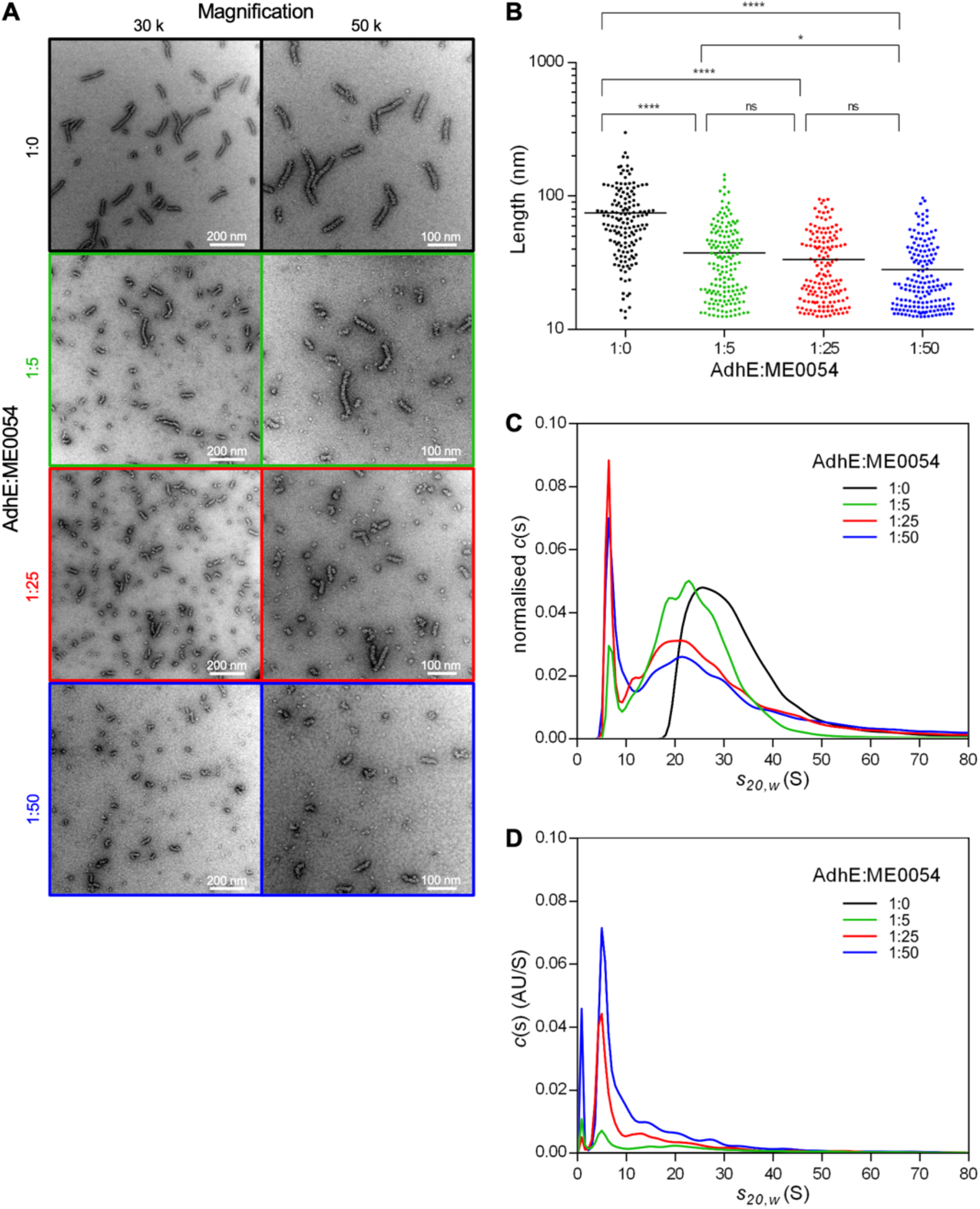
ME0054 disrupts AdhE spirosomes. AdhE fraction 12 containing long spirosomes was incubated for 1 h at 37°C with ME0054 at different (1:0, 1:5, 1:25 and 1:50) molar ratios. Throughout this figure black signifies an AdhE:ME0054 molar ratio of 1:0, green 1:5, red 1:25 and blue 1:50. The AdhE spirosome disruption produced by ME0054 was visualised by **(A)** negative stain EM and **(B)** the lengths of at least 100 spirosomes from each condition were measured using ImageJ software (21). SV-AUC was performed at 30 k rpm with absorbance wavelength set to **(C)** 280 or **(D)** 330 nm for observation of AdhE or ME0054, respectively. ns, not significant. * and **** indicate P < 0.05 and 0.0001 respectively from N ≥ 100 (ordinary one-way ANOVA test). The mean for each sample is represented by a horizontal black line.

### ME0054 disrupts AdhE spirosomes within 4 minutes

Computation of SAXS profiles for AdhE spirosome models (Fig. 1*D*) confirmed that it is possible to distinguish between and determine the dominant spirosome lengths in a given AdhE fraction based on SAXS data (Fig. 1*E*). We decided to use time resolved (TR) SAXS to explore the kinetics of spirosome disruption by ME0054. AdhE F12 was mixed with ME0054 at a molar ratio of 1:10 (AdhE:ME0054) and SAXS data recorded every 3 min for 60 min at 37°C (Fig. 4*A*). Due to limitations imposed by the beamline environment, it was not possible to acquire data at this temperature before 4 min. Comparison of the results obtained after 4 min at 37°C in the presence or absence of ME0054 clearly shows changes in the features at q = 0.083 Å^−1^ and q = 0.057 Å^−1^ (see e.g. Fig. 1*D, E*) consistent with a reduction in spirosome length produced by ME0054 (Fig. 4*A*). No obvious changes were detected at low angle (q < 0.025 Å^−1^). In the absence of compound, there were no changes in the SAXS profile over the observation period of 60 min (data not shown). Interestingly, after 4 min, changes to the profiles recorded in the presence of ME0054 became more subtle, suggesting that most of the effect of the compound was produced by that time. To try to follow the process of spirosome disruption at a higher time resolution, we decreased the temperature to 25°C, expecting that spirosome disruption might be slowed somewhat. To carry out the TR-SAXS experiments at 25°C, we used a novel microfluidic stopped-flow chip that allowed us to acquire data every 1 min from 0.06 s, when mixing of the AdhE with ME0054 was complete. This new device allowed us to minimise sample volumes, maximise the number of data recorded and significantly reduce dead time at the start of data acquisition. As expected, we managed to observe the evolution of key features in the SAXS profile in greater detail and over a longer period (59 min) as the result of the disruption of AdhE spirosomes by ME0054 at 25°C (Fig. 4*B*). Taken together these results confirm that at biologically relevant temperatures ME0054 disrupts AdhE spirosomes within 4 min but that the process of disruption continues for at least 55 min thereafter.

**Figure 4.**
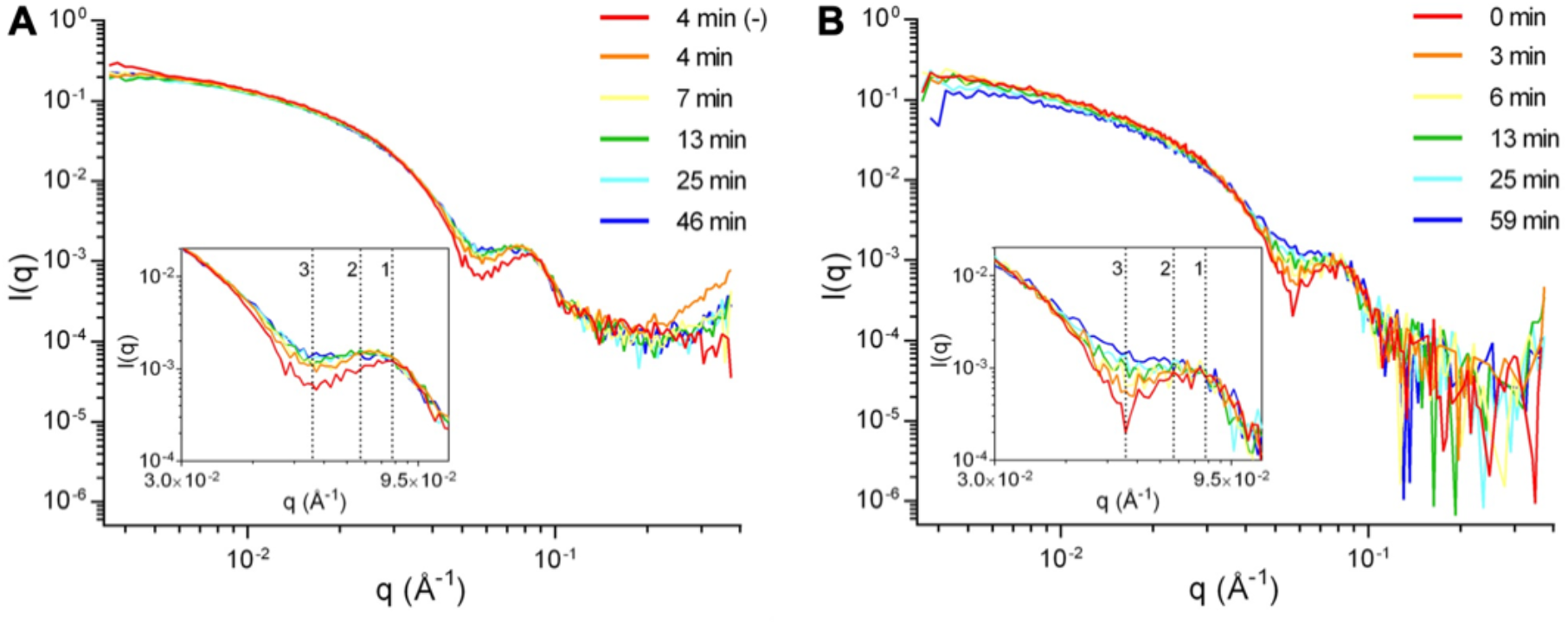
Kinetics of ME0054 by time-resolved (TR) SAXS. AdhE fraction 12 containing long spirosomes was mixed with ME0054 at a molar ratio of 1:10 (AdhE:ME0054) in **(A)** a quartz capillary at 37°C or **(B)** a microfluidic device at 25°C for the acquisition of SAXS data for 1 h every 3 or 1 min, respectively. The evolution of features at q = 0.083 Å^−1^ (feature 1) to q = 0.072 Å^−1^ (feature 2) together with a diminution of that q = 0.057 Å^−1^ (feature 3) is evident, consistent with a loss of the helical repeat.

## DISCUSSION

The bacterial protein AdhE presents a captivating enigma on multiple fronts. It forms spirosomes, crucial for its activity, yet the requirement for such a diverse range of spirosome lengths remains shrouded in mystery. This fundamental gap in understanding the structure-function relationship of AhdE has impeded our understanding of how this protein impacts bacterial metabolism.

AdhE is also of interest as it was identified as a target protein of the SA family of AV compounds. Evidence for this came from an affinity chromatography experiment that found the SA compounds bound numerous bacterial proteins including AdhE (7). Our prior investigations delved into the phenotypic outcomes associated with the deletion of the *adhE* gene in EHEC. Notably, we observed a robust suppression of the T3SS and a 20% increase in extracellular acetate levels. This phenotype is similar to that observed when this strain is exposed to SA compounds (8), implying that AdhE is one of the key targets of these compounds. However, how the SA compounds caused these phenotypes was unresolved and has spurred us to better understand the structure and function of AdhE.

In this work we replicated previous results showing by negative stain EM that purified AdhE with a 6-His tag fusion at the N-terminal is able to oligomerise, forming helicoidal filaments heterogeneous in length called spirosomes (Fig. 1*A*). Understanding why AdhE spirosomes are formed with different lengths seemed key to unravelling the basis to how this protein functions, and, by implication, how we can inhibit it.

AdhE is a bifunctional aldehyde-alcohol dehydrogenase enzyme, highly conserved in bacteria, fungi, algae and protozoa. The N-terminal domain is an aldehyde dehydrogenase (ALDH) that converts acetyl-CoA to acetaldehyde, a toxic intermediate. The acetaldehyde is converted to ethanol by the AdhE C-terminal alcohol dehydrogenase (ADH) domain (9). The N-terminal ALDH is connected to the C-terminal ADH by a linker. The AdhE dimer is formed by the interaction of two N-terminal domains with the formation of continual β-sheet between the three-stranded β-sheet in the linker and the β-sheet within the ALDH domain from the other molecule (9). Then, the tetramer is formed by the interaction of two ADH domains forming one helical pitch. Thus, the spirosome structure can have different lengths depending on the number of repetitions of the helical unit based on the number of monomers incorporated. Therefore, a long spirosome contains a higher number of ALDH and ADH domains compared with a short one and consequently could be enzymatically more efficient. Interestingly, the two activities are topologically separated in the spirosome, suggesting that these filaments work as substrate channels, avoiding the exposure of the bacterial cytoplasm to the toxic aldehyde intermediate. The high-resolution cryo-EM structure of one-and-a-half helical turns showed that the ALDH domains are located towards the outer surface of the helical structure, whereas the ADH domains point towards the inner surface (9). In addition, it is important to highlight the fact that AdhE is bidirectional, and therefore also able to convert ethanol to acetyl-CoA. Another explanation for variable spirosome length could be that filaments with different accessibilities to the ALDH and ADH domains, due to the spirosome length, would be more/less efficient in one direction of the reaction thus, the differences in spirosome lengths would be necessary to control AdhE bidirectionality. To explore these options, in this work AdhE spirosomes were fractionated by length (Fig. 1*B*) by SEC to compare the enzymatic efficiency of long versus short filaments in the two directions of the reaction (Fig. 2*B, C*).

In size exclusion chromatography (SEC), proteins elute from the column based on their size (20), species with higher hydrodynamic volume eluting earlier. Because AdhE spirosomes are heterogeneous in length, we expected the chromatogram to comprise multiple peaks and, in this way, fractions to be homogeneous in spirosome size. However, AdhE eluted in a single asymmetric peak (Fig. 1*B*), so fractions were collected during the elution and analysed for homogeneity or otherwise. SV-AUC confirmed that AdhE was only partially fractionated (Fig. 1*C*). Because the protein was fractionated by SEC, these findings suggest that spirosome formation could be a dynamic process. The spirosomes could be broadly fractionated based on hydrodynamic volume but perhaps re-equlibrate to broader oligomeric distributions after elution. Further exploration should confirm this possibility.

To understand why AdhE forms spirosomes with different lengths, we formulated two plausible hypotheses: (a) longer spirosomes are enzymatically more efficient or (b) spirosome length drives the direction of the enzymatic reaction. Once fractions containing a main population of long (F12) or short (F24) spirosomes were identified, enzymatic assays were carried out to compare their activities both in the forward and reverse reactions (Fig. 2*A*-C). The spirosome length did not have any impact in the forward reaction (Fig. 2*B*). However, short spirosomes were more efficient during the reverse reaction (Fig. 2*C*). The ALDH and ADH domain configuration in the spirosome helical structure could explain why short spirosomes are more efficient in the reverse reaction. Thus, our findings refute hypothesis (a) and support hypothesis (b). We propose the following model to explain the relationship between spirosome length and enzymatic reaction direction (Fig. 5). When the filament is short, the ADH domains from the inner surface of the filament could be more accessible for the ethanol substrate than when consecutive ADH domains are within the filament, whereas the disposition of the ALDH domains towards the spirosome outer surface makes them always accessible to the acetyl-CoA substrate, independent of spirosome length. The observation that the conversion of acetyl-CoA to ethanol (forward reaction) has the same efficiency independent of the spirosome length (Fig. 5) could be relevant for industrial biofuel production. Our results demonstrate that control of spirosome length would be unnecessary for ethanol production by AdhE, simplifying the process and making it possible for unfractionated protein to be used as the biological catalyst. Nevertheless, the fact that short spirosomes are more efficient in the conversion of ethanol to acetyl-CoA allows us to envision at least three different scenarios: short, long or a mix of lengths as intermediate situation (Fig. 5). In a situation where only short spirosomes are present, the efficiency of the reverse reaction would be very high, reducing the levels of ethanol and accumulating acetyl-CoA in the bacterial cytoplasm. AdhE spirosome formation would be necessary to decrease the efficiency of the reverse reaction (Fig. 5). Because the efficiency of the forward reaction is constant, in an intermediate situation with a mix of short and long spirosomes both directions of the reaction are balanced (Fig. 5). Taken together, these results suggest that AdhE spirosome formation, coupled with dynamic disassembly of the filaments, is necessary to drive the enzymatic bidirectionality of AdhE.

**Figure 5.**
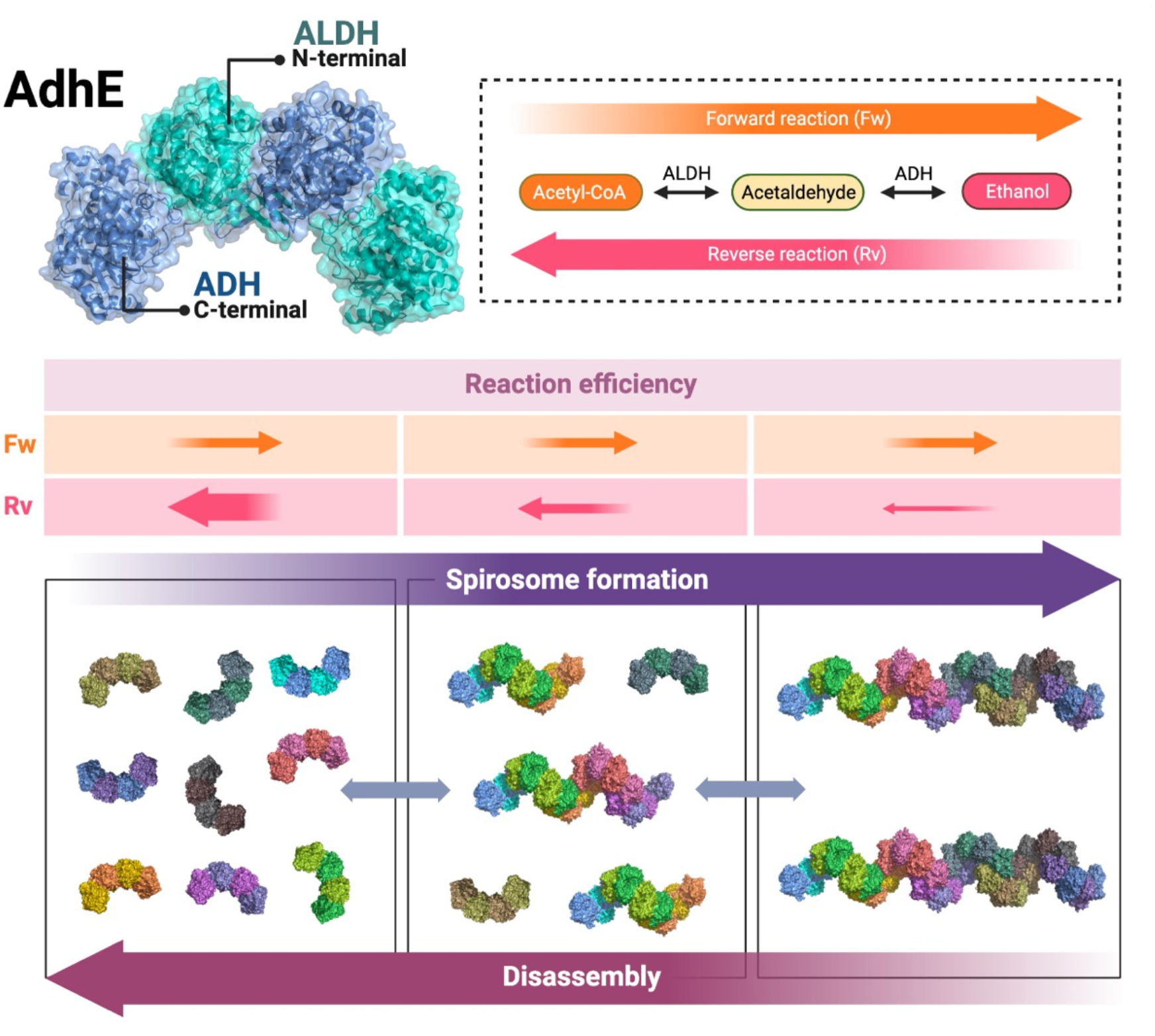
Model explaining that AdhE needs to form spirosomes to balance the direction of the enzymatic reactions. **Top**: AdhE dimer model and schematic representation of the AdhE enzymatic activity in the forward (Fw) and reverse (Rv) reactions. **Bottom**: 3 scenarios showing short, long or a mix of spirosome lengths during AdhE spirosome formation or disassembly. **Middle**: The forward (Fw) reaction efficiency is constant in the different scenarios whereas short spirosomes are more efficient during the reverse (Rv) reaction. Created with BioRender.com.

AdhE has been mooted as potential target for the SA compounds (8). Here for first time, we formally confirm this with the observation of a direct effect of the SA compound named ME0054 on AdhE spirosomes. The enzymatic activities of fractions containing a main population of long (F12) or short (F24) spirosomes were tested in the presence of ME0054 (Fig. 2*B, C*). No effect on activity was observed in the forward reaction in the presence of compound (Fig. 2*B*). However, the spirosome activity increased in the presence of ME0054 during the reverse reaction (Fig. 2*C*). Because short spirosomes are more efficient in the reverse reaction this result suggests that ME0054 could be disrupting the spirosomes and therefore enhancing the efficiency of the reverse reaction. To confirm this assumption, AdhE F12 comprising a main population of long spirosomes was incubated with ME0054 at different protein:compound molar ratios. Negative stain EM combined with SV-AUC showed that ME0054 disrupted AdhE spirosomes in a concentration dependent manner (Fig. 3). Even though the mean spirosome length decreased when the concentration of compound increased (Fig. 3*B*), ordinary one-way ANOVA tests revealed that the differences in length were statistically significant only when the control without ME0054 was compared with different compound concentrations, whereas there were no significant differences between the mean spirosome lengths observed at different ME0054 concentrations (Fig. 3*B*). However, the smallest molecules could not be detected by negative stain EM due to the magnification scale. Instead, to quantify all molecules present in the sample, SV-AUC was performed by measuring the absorbance at 280 nm to follow AdhE spirosomes. We observed how the sedimentation coefficient (s_20,w_) distribution for the sample without ME0054 was displaced to the left and split in two in the presence of compound (Fig. 3*C*). The decrease in s_20,w_ directly correlates with a reduction in the protein size, confirming the observation from negative stain EM. In the presence of compound, we can observe how the peak with s_20,w_ ∼ 20 S decreased, whereas a peak with s_20,w_ ∼ 5 S increased. The result confirmed that by increasing the compound concentration there was a greater disruption of spirosomes and species with higher s_20,w_ were converted to species with lower s_20,w_.

However, the precise mechanism of action of ME0054 on AdhE remains unknown. To at least examine whether the compound is bound to the spirosome or simply disrupts and then dissociates from it, SV-AUC data were acquired at a wavelength of 330 nm (Fig. 3*D*). At this wavelength ME0054, but not AdhE, absorbs. The molecular weight of ME0054 is 272 Da, as such it would be expected to sediment with s_20,w_ ∼ 0.2 S (and indeed a peak at very low s_20,w_ is evident in Fig. 3*D*). The co-sedimentation of the compound with AdhE both at high and low s_20,w_ suggests that ME0054 binds to both long and short spirosomes. Finally, the effect of ME0054 on long spirosomes (F12) was tested at 37°C and 25°C by TR-SAXS (Fig. 4). The compound acted more rapidly at 37°C than at 25°C, but was still active at 25°C.

This result is promising in the context of using ME0054 to treat EHEC infections in humans, where the average temperature is 37°C.

AdhE spirosomes disrupted by ME0054 are shorter and thus more efficient in the reverse reaction (Fig. 2C). Therefore, addition of ME0054 enhances the consumption of ethanol and increases the accumulation of acetyl-CoA in the bacterial cytoplasm. Intriguingly, previous studies have shown that deletion of *adhE* results in a 20% increase in acetate secretion compared with the wild-type parent (8). Such an increase would result in a 9 mM increase in the intracellular acetate pool, a substantial perturbation. Hence our finding that ME0054 disrupts AdhE spirosomes, thereby favouring accumulation of acetyl-CoA, may provide an explanation for the SA compounds mirroring some of the phenotypes seen by deletion of *adhE*. However, even though in this work AdhE has been validated as a protein target of ME0054, we cannot discard the possibility that ME0054 may interact with other proteins to induce the observed bacterial phenotypes.

Taken together, these results confirm that AdhE is indeed a protein target for ME0054 and that the compound acts by disrupting the AdhE spirosomes. This step forward in our understanding of how the compound works will help in the design of more specific and targeted inhibitors of AdhE that can act as alternatives to traditional antibiotics.

## MATERIALS AND METHODS

### Protein purification and size exclusion chromatography (SEC)

The *adhE* gene from *Escherichia coli* K12 was cloned into a pET28a vector for expression with an N-terminal 6-His tag. Plasmid expressing recombinant 6-His-AdhE protein was transformed into BL21(DE3) competent *E. coli* cells. A single colony was used to inoculate an overnight culture at 37°C with shaking at 200 rpm prior to back-diluting into 4 L fresh LB supplemented with kanamycin (50 μg ml^-1^) and cultured until OD_600 nm_ = 0.8 was reached. AdhE overexpression was induced by addition of IPTG to a final concentration of 0.5 mM and cultures were grown overnight at 18°C. Cells were harvested by centrifugation for 10 min at 5,000 rpm and the supernatant removed. The cell pellet was resuspended in buffer A (50 mM Tris-HCl pH 8.0, 500 mM NaCl, 5% (v/v) glycerol) containing 5 mM imidazole, lysozyme, EDTA-free protease inhibitor cocktail (Promega) and DNase. Cells were lysed by sonication on ice (1 s on, 1 s off for 6 min) and then centrifuged for 50 min at 4°C at 18,000 rpm. After centrifugation, the 6-His-AdhE protein was found in the soluble fraction. The supernatant was removed and filtered through a 0.22 μM filter and loaded onto a HisTrap High Performance column (GE Healthcare) equilibrated with buffer A containing 5 mM imidazole. The column was washed with a linear gradient (5-75 mM) of imidazole in buffer A and 6-His-AdhE was then eluted in 50 ml buffer A with 300 mM imidazole. The 50 ml containing 6-His-AdhE was concentrated to a final volume of 10.5 ml using a Vivaspin 20 centrifugal concentrator (Sartorius) with a molecular weight cut-off of 30 kDa and then dialysed against buffer A. A single aliquot (0.5 ml) of unfractionated 6-His-AdhE in buffer A was stored at -20°C.

For protein fractionation by size exclusion chromatography (SEC), the remaining 10 ml of concentrated 6-His-AdhE in buffer A were loaded onto a HiLoad 26/600 Superdex 200 pg column (GE Healthcare) equilibrated with buffer A. 2 ml fractions were eluted with buffer A and stored at -20°C. Six fractions were selected as representative fractions (F12, 14, 16, 18, 21 and 24) for subsequent analyses.

### Negative stain electron microscopy (EM) and spirosome length quantification

For visualisation of unfractionated 6-His-AdhE, the protein was diluted in buffer B (50 mM Tris-HCl pH 8.0, 500 mM NaCl) to a final concentration of 10 μg ml^-1^. To observe the effect of ME0054 compound on 6-His-AdhE, ME0054 stock solution was prepared in dimethyl sulphoxide (DMSO), aliquoted, and stored at -20°C in the dark. Fraction 12 (F12) from SEC (2.8 μM) was incubated with ME0054 for 1 h at 37°C at different AdhE:ME0054 molar ratios (1:5, 1:25 and 1:50). As a control without compound (1:0), F12 was incubated with the same amount of DMSO (0.14%) used in the sample with the highest compound concentration (i.e. AdhE:ME0054 = 1:50) to confirm that any observed effect is not due to the DMSO. After incubation, the samples were diluted 30-fold in buffer B. Drops (5 μl) of each diluted sample were applied to glow-discharged carbon coated Cu (400 mesh) grids and incubated for 5 min. The grids were washed three times with water and incubated with 2% (w/v) uranyl acetate for 5 min for negative staining. Excess uranyl acetate was removed by filter paper and the grids dried. Prepared grids were analysed using a JEM-1400Flash transmission electron microscope (TEM) (JEOL) running at 80 keV, images captured using a CDD camera and TEM Center software version 1.7.26.3016 (JEOL).

For the quantification of spirosome dimensions, the lengths of at least 100 spirosomes from each condition were measured using ImageJ software (21) and analysed using ordinary one-way ANOVA tests.

### Small angle X-ray scattering (SAXS)

SAXS was carried out on beamline B21 of the Diamond Light Source synchrotron facility (Didcot, UK). Data were recorded at 13 keV, at a sample-detector distance of 3.6 m using an Eiger 4M detector (Dectris, Switzerland). For batch mode measurements, 40 μl of AdhE representative fractions (final concentration 1.0 mg ml^-1^) were loaded into a 96-well plate, before being sequentially injected into a quartz capillary by the BioSAXS automatic sample changer (ARINAX). Data were acquired at 6°C and processed with ScÅtter (http://www.bioisis.net) as previously described (9). SAXS profiles for AdhE spirosome models were computed with the FoXS server (14, 15), which was also used to fit models against experimental data (Fig. S2, Table S1).

### Sedimentation velocity analytical ultracentrifugation (SV-AUC)

SV-AUC was performed using a Beckman Coulter XL-I analytical ultracentrifuge equipped with an An-50 Ti eight-hole rotor as previously described (9). 300-360 μl samples (AdhE final concentration 0.28 mg ml^-1^ in buffer A) were loaded into 12 mm pathlength charcoal-filled epon double-sector centrepieces, sandwiched between two sapphire windows and equilibrated at 4°C in vacuum for 6 h before running at 30 k rpm. To examine the effect of ME0054, AdhE fractions were incubated with ME0054 at different AdhE:ME0054 molar ratios (1:0, 1:5, 1:10, 1:25 and 1:50, in buffer B) during 1 h at 37°C prior to loading. Absorbance data were recorded in continuous mode between radial positions of 5.65 and 7.25 cm, with a radial resolution of 0.005 cm and a nominal time interval of 7 min. Wavelengths of 280 nm or 330 nm were used to follow AdhE protein or ME0054 compound, respectively. Data were analysed with the program SEDFIT (22) using a continuous c(s) model and normalised by area with GUSSI (23). The partial specific volume, buffer density and viscosity were calculated using SEDNTERP (24) (Table S2).

### Enzymatic assays

To determine the enzymatic activity of AdhE fractions (F12 or 24) in the forward and reverse reactions, the consumption or production of NADH, respectively, was measured at a wavelength of 340 nm using a FLUOstar Omega microplate reader (BMG Labtech). All assays were performed at 37°C and the total volume was 100 μl. All reagents were prepared in 50 mM Tris-HCl pH 8.0 buffer. For measuring the conversion of acetyl-CoA to ethanol (forward reaction), the assays were performed with reaction mixtures containing 50 mM Tris-HCl pH 8.0, 250 mM NaCl, 20 μM FeSO_4_, 200 μM acetyl-CoA, and 250 μM NADH with 60 μg of AdhE with or without ME0054 at a molar ratio of 1:10 (AdhE:ME0054). The conversion of ethanol to acetyl-CoA (reverse reaction) was measured for reaction mixtures containing 50 mM Tris-HCl pH 8.0, 250 mM NaCl, 20 μM FeSO_4_, 200 μM CoASH, 200 mM ethanol, and 500 μM NAD^+^ with 6 μg of AdhE with or without ME0054 at a molar ratio of 1:10 (AdhE:ME0054).

### Time-resolved (TR) SAXS

TR-SAXS was performed on beamline ID02 at the European Synchrotron Radiation Facility (ESRF) (Grenoble, France). For the acquisition of data at 37°C, 50 μl of AdhE F12 (final concentration 3.0 mg ml^-1^) with or without ME0054 at a molar ratio of 1:10 (AdhE:ME0054) were placed in a quartz capillary of external diameter 1.1 mm. Nine frames of data from 0.1 s exposures were collected every 3 min over a period of 60 min. The first measurement was taken 4 min after the sample was placed in the capillary. For the acquisition of data at 25°C, a novel X-ray transparent (Kapton window) microfluidic stopped-flow chip (developed at University of Leeds) was used. One syringe was loaded with 1 ml of AdhE F12, the second contained 0.5 ml ME0054 (or DMSO as negative control). The first shot (to fill the tubing between the syringes and the chip) delivered 140 μl from each syringe. Consecutive shots delivered 37.8 μl from the syringe containing AdhE and 4.2 μl from the syringe containing ME0054 to generate a molar ratio of 1:10 (AdhE:ME0054) with AdhE at a final concentration of 3.0 mg ml^-1^. Sixty frames of data from 0.05 s exposures were collected every 1 min over a period of 59 min. The acquisition was performed in triplicate and the average results plotted.

## ACKNOWLEDGMENTS

Special thanks to our long-term collaborator, Mikael Elofsson (University of Umeå, Sweden), who provided the SA compounds. We thank Margaret Mullin from the Electron Microscopy Facility at the University of Glasgow for help with negative stain EM data collection. We are grateful to Diamond Light Source for the SAXS beamtime at B21 beamline and to the European Synchrotron Radiation Facility for the TR-SAXS beamtime at ID02 beamline and we thank the beamline scientists, Nathan Cowieson, Charlotte Edwards-Gayle, Katsuaki Inoue, Nikul Khunti and Lauren Matthews, for excellent scientific support. The research conducted by E.S., A.J.R. and O.B. was supported by the BBSRC, project grant reference BB/V009494/1.

## SUPPLEMENTARY FIGURES AND TABLES

**Figure S1.**
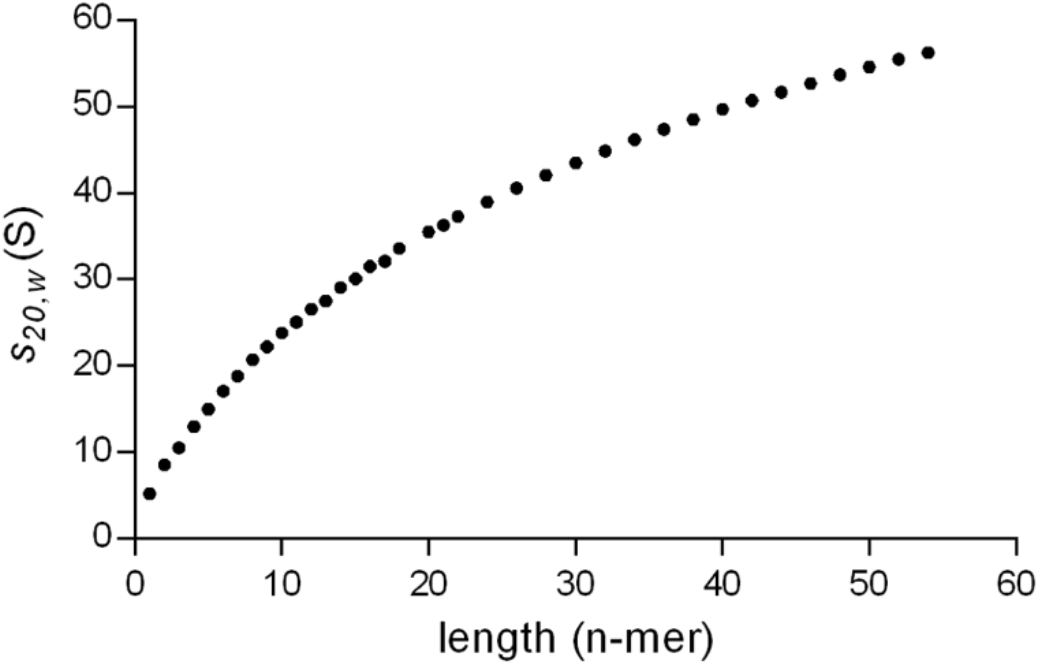
Representation of sedimentation coefficients (s_20,w_) computed for AdhE spirosomes of different lengths. The atomic resolution structure of AdhE (PDB ID 6AHC [6]) was used to generate spirosome models of varying lengths. AtoB (13) was used to compute grid-based bead models for which hydrodynamic calculations (including the sedimentation coefficient) were undertaken using SOMO (14,15), permitting the construction of a plot of s_20,w_ versus oligomer length.

**Figure S2.**
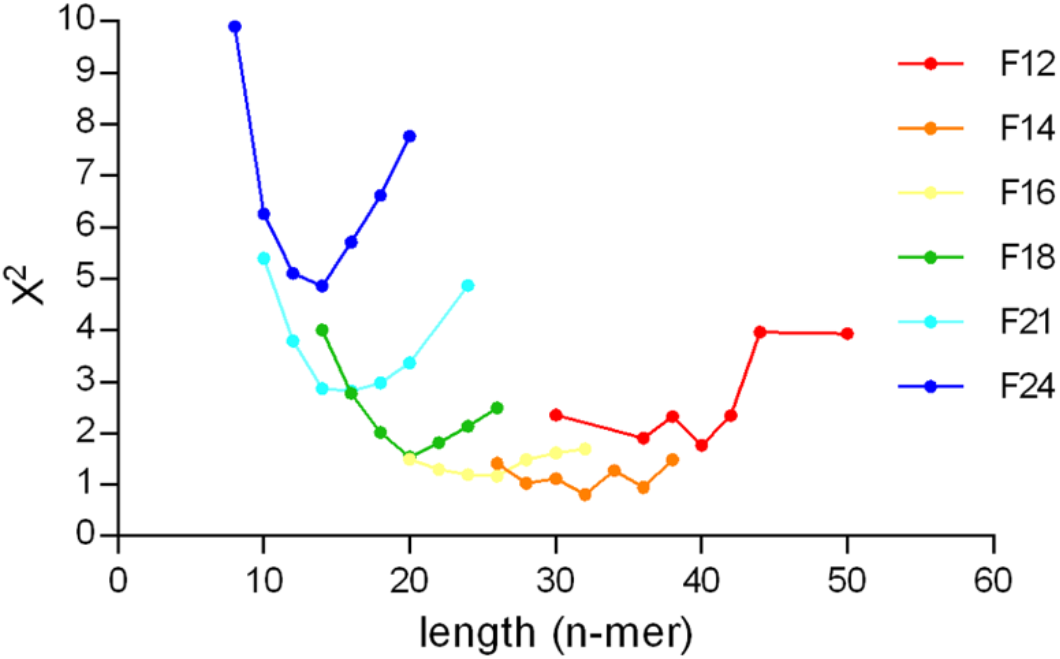
Graphical representation of fits to SAXS data for AdhE representative fractions by AdhE spirosome models of different oligomeric lengths. The goodness of fit (represented by the χ^2^ parameter) is plotted as a function of the number of monomers in the spirosome model. The atypical dependence of χ^2^ observed for the fit to F12 SAXS data by the 50-mer model (right-hand-most red datum) likely arises because any differences between the computed and experimental SAXS curve will occur at such low angles that they do not contribute to an increase in χ^2^ (compared with the value obtained for the fit to F12 SAXS data by the 44-mer model). The best fitting models, with the lowest χ^2^ for each fraction, are summarised in Table S1.

**Table S1.** Best fits to SAXS data for AdhE representative fractions by AdhE spirosome models of different lengths. The goodness of fit is represented by the χ^2^ parameter. The best performing models were determined from Fig. S2.

| Fraction | $\chi^2$ | Model (n-mer) |
| --- | --- | --- |
| F12 | 1.76 | 40 |
| F14 | 0.81 | 32 |
| F16 | 1.17 | 26 |
| F18 | 1.54 | 20 |
| F21 | 2.82 | 16 |
| F24 | 4.86 | 14 |

**Table S2.**
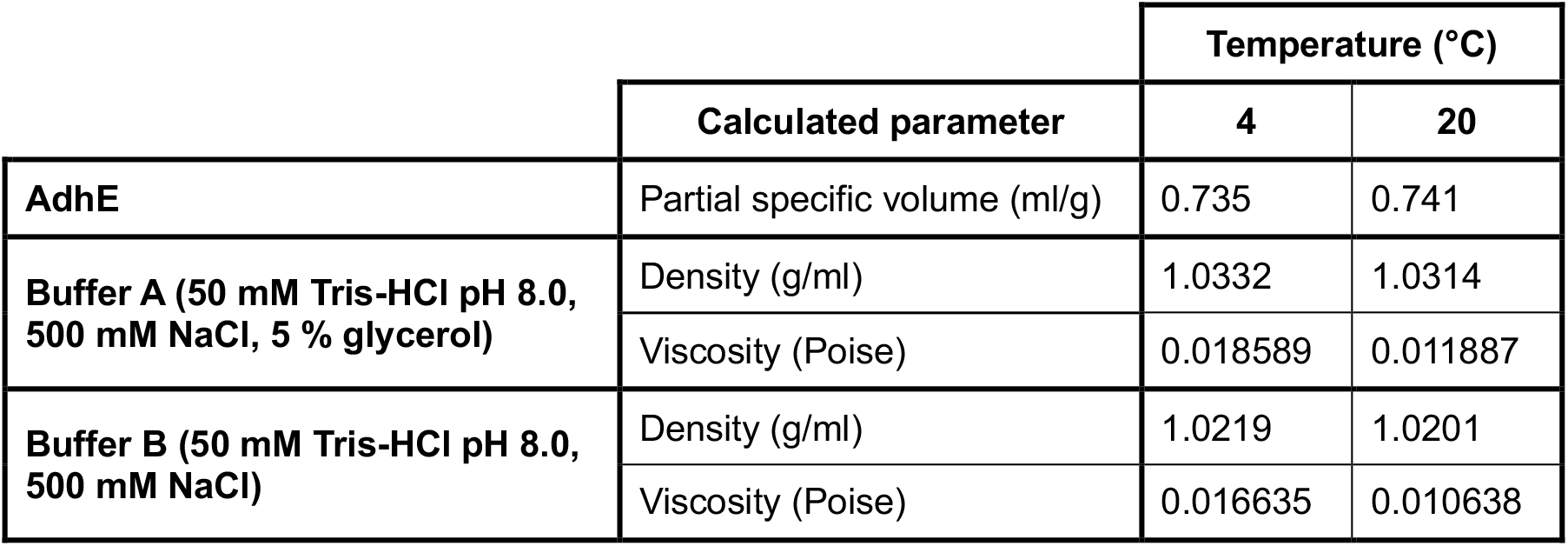
Partial specific volume, buffer density and viscosity calculated using SEDNTERP (24).

